# Lytic Replication and Reactivation from B Cells Is Not Required for Maintaining Gammaherpesvirus Latency *in vivo*

**DOI:** 10.1101/2021.08.20.457181

**Authors:** Arundhati Gupta, Shana M. Owens, Darby G. Oldenburg, Douglas W. White, J. Craig Forrest

## Abstract

Gammaherpesviruses (GHVs) are lymphotropic tumor viruses with a biphasic infectious cycle. Lytic replication at the primary site of infection is necessary for GHVs to spread throughout the host and establish latency in distal sites. Dissemination is mediated by infected B cells that traffic hematogenously from draining lymph nodes to peripheral lymphoid organs, such as the spleen. B cells serve as the major reservoir for viral latency, and it is hypothesized that periodic reactivation from latently infected B cells contributes to maintaining long-term chronic infection. While fundamentally important to an understanding of GHV biology, aspects of B cell infection in latency establishment and maintenance are incompletely defined, especially roles for lytic replication and reactivation in this cell type. To address this knowledge gap and overcome limitations of replication-defective viruses, we generated a recombinant murine gammaherpesvirus 68 (MHV68) in which *ORF50*, the gene that encodes the essential immediate-early replication and transcription activator protein (RTA), was flanked by *loxP* sites to enable conditional ablation of lytic replication by *ORF50* deletion in cells that express Cre recombinase. Following infection of mice that encode Cre in B cells with this virus, splenomegaly and viral reactivation from splenocytes were significantly reduced, however the number of latently infected splenocytes was equivalent to WT MHV68. Despite *ORF50* deletion, MHV68 latency was maintained over time in spleens of mice at levels approximating WT, reactivation-competent MHV68. Stimulation of polyclonal B cell activation and proliferation by treating mice with lipopolysaccharide (LPS), which promotes MHV68 reactivation *ex vivo*, yielded equivalent increases in the number of latently infected cells for both *ORF50*-deleted and WT MHV68, even when mice were simultaneously treated with the antiviral drug cidofovir. Together, these data demonstrate that lytic replication in B cells is not required for MHV68 latency establishment and maintenance and further indicate that B cell proliferation, and not reactivation *per se*, is a major mechanism for maintaining latent viral genomes in the host.

**IMPORTANCE:** Gammaherpesviruses establish lifelong chronic infections in cells of the immune system and place infected hosts at risk for developing lymphomas and other diseases. It is hypothesized that gammaherpesviruses must initiate acute infection in these cells to establish and maintain long-term infection, but this has not been directly tested. We report here the use of a viral genetic system that allows for cell-type-specific deletion of a viral gene that is essential for replication and reactivation. We employ this system in an *in vivo* model to reveal that viral replication is not required to initiate or maintain infection within immune cells.

## INTRODUCTION

Herpesviruses are large, enveloped viruses that contain a double-strand DNA genome within a protein capsid and establish life-long chronic infections (1). The gammaherpesvirus (GHV) subfamily includes the human pathogens, Epstein-Barr virus (EBV) and Kaposi sarcoma herpesvirus (KSHV), which infect a large percentage of the adult population worldwide. GHV infections do not typically result in severe disease in healthy individuals, but can cause a variety of cancers, especially in immunocompromised patients (2–9). EBV is the causative agent of infectious mononucleosis and is associated with Burkitt lymphoma, nasopharyngeal carcinoma, post-transplant lymphoproliferative disorder, Hodgkin lymphoma and other cancers in chronically infected persons (3, 8, 10–12). KSHV is the etiologic agent of Kaposi sarcoma, an AIDS-defining malignancy, as well as multicentric Castleman disease and primary effusion lymphoma (2, 4–6, 9, 13–15). These diseases are epidemiologically and economically impactful, and laboratory-based pathogenesis studies are important.

Like all herpesvirus infections, the GHV infection cycle is biphasic. The productive replication phase, a process known as lytic replication, occurs following primary infection of a host. Here, viral gene products are expressed in a regulated cascade and infectious viral particles are produced (16). The second phase of infection is chronic and persists for the life of the infected host. This phase, known as latency, is characterized by minimal expression of viral genes as the virus maintains the genome as a circular episome within the host-cell nucleus (17, 18). Proteins expressed by the virus in latency facilitate maintenance of the viral genome and promote cellular proliferation and survival (19–23). Upon receiving an appropriate stimulus, a latently infected cell can reactivate and re-initiate the lytic cycle.

Since chronic GHV infection can lead to severe disease, understanding the fundamental mechanisms of viral infection and maintenance of chronic infection is essential. However, human GHVs exhibit a very narrow host range and do not readily infect laboratory animals (17, 24–26). Murine gammaherpesvirus 68 (MHV68) is a natural pathogen of murid rodents that provides a highly tractable small-animal model for studying GHV infection and latency *in vivo*. MHV68 is a close genetic relative of the human GHVs; genes essential for viral replication are largely conserved amongst the GHVs, and there is overlap in the host RNAs targeted by EBV, KSHV, and MHV68 non-coding RNAs (27, 28). Several MHV68 and KSHV latency gene products also are conserved and functionally interchangeable (29–31). MHV68 pathogenesis recapitulates key aspects of human GHV infections, targeting B cells as the preferred reservoir for latency and promoting development of lymphomas and lymphoproliferative diseases following infection (24).

The MHV68 mouse model has shaped our understanding of GHV infection, dissemination, and persistence in a host. Following intranasal inoculation of mice, MHV68 undergoes lytic replication in the respiratory epithelium (32). This enables MHV68 to infect tissue-resident and infiltrating immune cells, cross the epithelial barrier, and traffic to draining lymph nodes (33–36). MHV68 is thought to expand in the lymph node prior to disseminating to distal sites, such as the spleen (35, 37, 38). In addition to B cells, MHV68 establishes latency in macrophages, dendritic cells, epithelial cells, and endothelial cells (39), however several lines of evidence indicate that B cells play an indispensable role in MHV68 trafficking to the spleen.

For example, MHV68 does not establish latency in the spleen following intranasal inoculation of B cell-deficient μMT mice, but dissemination to the spleen is restored if B cells are adoptively transferred into these animals prior to infection (40, 41). Experiments in which *ORF73* and *M2*, latency-associated genes, were conditionally deleted in B cells demonstrated accumulation of virus in draining lymph nodes after IN inoculation, but reduced blood infection and a drastic reduction in latency in the spleen, indicating the importance of the B cell population in viral dissemination (33, 42). Interestingly, replication defective viruses also either fail to disseminate after IN inoculation or exhibit severe attenuation in the spleen (43–45), suggesting that lytic replication also plays a pivotal role in seeding distal organs with latent virus. Further, despite the clear importance of B cells and lytic replication in systemic MHV68 dissemination and latency, it is not known if lytic replication *within* the B cell compartment is required for trafficking to and establishing latency in distal cellular reservoirs.

Additionally, the mechanisms used by GHVs to maintain latency within specific cellular populations is not completely defined. Within the spleen, latently infected B cells are the predominant reactivation competent cell type in MHV68-infected mice, and will reactivate if explanted, especially if cells are treated with B cell-activating stimuli (46). Clearly, reactivation can provide a means for herpesviruses to “get out” and infect new hosts. However, it is also suggested that periodic “homeostatic reactivation” contributes to the long-term maintenance of latency within a host. This hypothesis is mostly based on observations suggesting that reactivation occurs in splenic B cells and macrophages during latency. For instance, adoptive transfer of B cells from mice latently infected with MHV68 into naïve mice leads to infection of B cells within recipient mice (47). Further, treating infected mice with TLR agonists, such as LPS, stimulates GHV reactivation from B cells and increases the number of infected splenocytes *in vivo* (48, 49). However, whether periodic reactivation, plays a major role in maintaining long-term latency *in vivo*, especially in immune competent animals, is not completely understood. Another viable hypothesis is that B cell proliferation due to non-GHV-related infections or other activating stimuli is sufficient to maintain the B cell latency reservoir. This has not been directly tested.

In experiments described here, we sought to determine whether GHVs require lytic replication within B cells in order to traffic systemically, establish, and maintain long-term latency. Traditional loss-of-function mutations in MHV68 lytic genes are valuable tools for defining functions of viral gene products during acute viral replication at the site of inoculation, but they are not suitable tools to answer questions pertaining to cell-specific requirements for essential lytic-cycle proteins or defining downstream roles for such proteins in viral pathogenesis (45, 50). To address this limitation, we generated an MHV68 recombinant virus that encodes *loxP* sites flanking a gene essential for lytic replication, *ORF50*. Utilizing cell-type-specific, Cre-mediated deletion of this gene, we define B cell-specific roles for lytic replication and reactivation in chronic GHV infection (51).

## RESULTS

### *Development and validation of a conditional* ORF50 *mutant*

The findings that (i) adoptive transfer of B cells into B cell-deficient mice restores trafficking to and latency establishment in the spleen after IN inoculation and (ii) that deletion of *ORF73*, the gene that encodes mLANA, specifically in B cells prevented trafficking from mediastinal LNs to the spleen demonstrate that B cells are critical for MHV68 to colonize the mouse spleen after IN inoculation. Replication defective viruses, such as those with null mutations in *ORF50* or *ORF31*, essential genes that respectively encode the replication and transcription activator (RTA) or a homolog of EBV BDLF4, fail to establish latency in the spleen following IN inoculation, suggesting the importance of lytic replication to latency establishment (43, 44, 52). However, whether lytic replication in the B cell compartment is necessary for viral dissemination to distal sites of latency is not known.

To address this question, we generated an MHV68 conditional mutant that can be rendered incapable of lytic replication by deletion of *ORF50* in a tissue-specific manner. We used a recombineering approach to insert *loxP* sites flanking the second exon of the *ORF50* gene in the MHV68 BAC to enable deletion of *ORF50* by Cre recombinase (O50.loxP MHV68) (**Fig. 1A**). To verify that *loxP*-flanked (floxed) *ORF50* was efficiently deleted from O50.loxP MHV68, we infected normal or Cre-expressing Vero cells with either WT MHV68 or O50.loxP and evaluated *ORF50* deletion by PCR (**Fig. 1B**). While intact *ORF50* was readily detected in Vero cells infected with O50.loxP, only the deletion product was detected in Vero-Cre cells infected with the conditional mutants. The intact locus was detected in both cell types following infection with WT MHV68. The *ORF73* gene was examined by PCR for off-target effects and found to be unaffected in all viruses tested. These finding demonstrate that floxed *ORF50* is deleted following infection of cells that express *Cre*.

**Fig 1.**
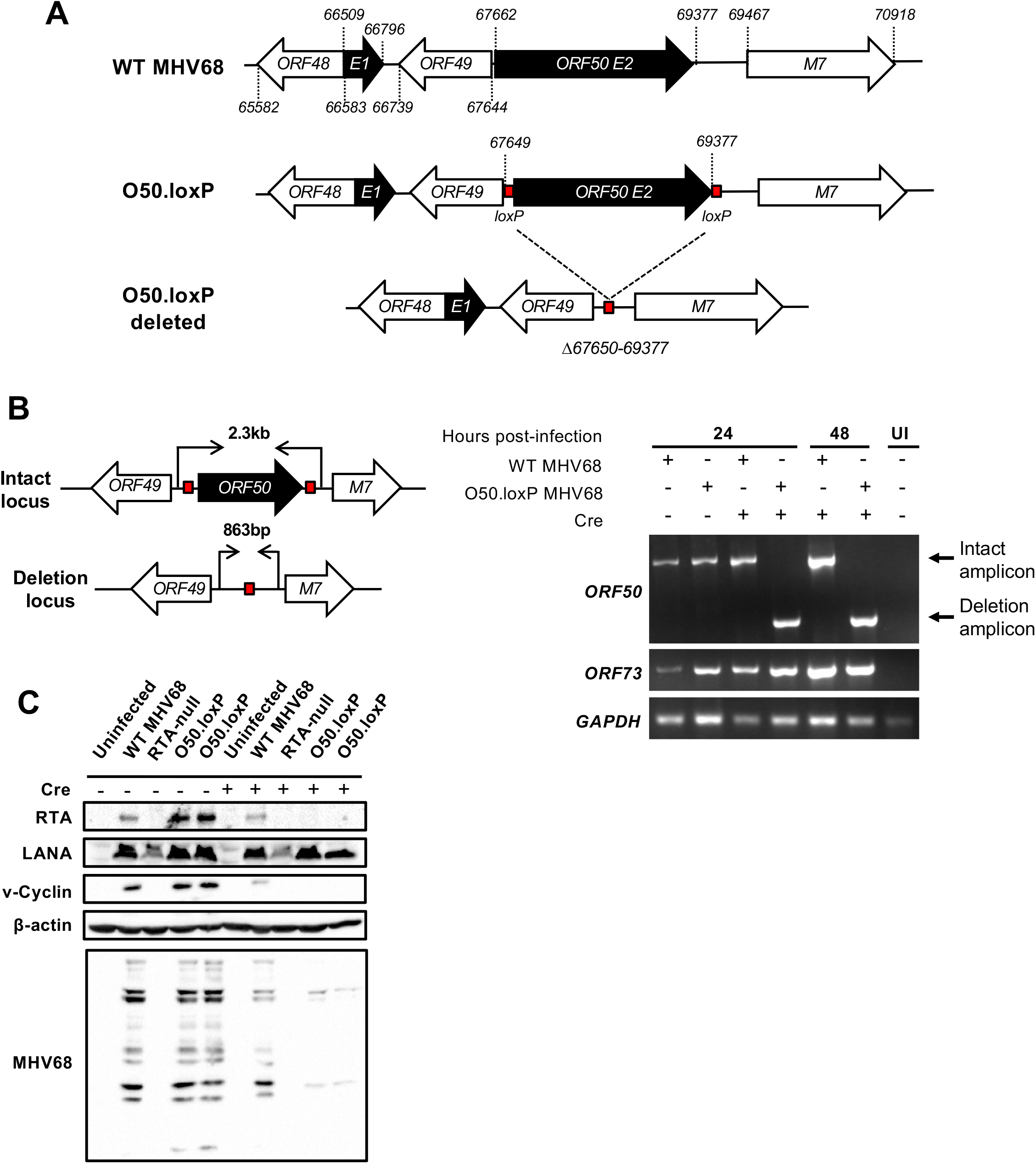
Derivation and validation of 50.loxP MHV68. (A) Schematic depicting the insertion of *loxP* sites flanking *ORF50* in the genome and its deletion after Cre-mediated recombination*. loxP* sites were inserted in the *ORF50* exon 2 (E2) after nucleotides *67649* and *69377* in the MHV68 genome. Cre-mediated recombination removes the *ORF50* E2 coding region (Δ*67650-69377*). (B) Vero cells or Vero-Cre cells were infected with the indicated virus at an MOI of 5 PFU/cell. Total DNA was isolated at 24 or 48 hours post-infection, and PCR was performed as illustrated in the schematic to detect the intact or deleted *ORF50* locus or the distal *ORF73* locus as a control. (C) 3T3 fibroblasts that encode Cre-ERT2 were treated with vehicle (−) or 4-hydroxytamoxifen (+) to induce Cre activity 24 h prior to infection. Treated cells were infected with the indicated viruses at an MOI of 0.5 PFU/cell. Cells were lysed on day 4 post-infection, and proteins were resolved by SDS-PAGE. Immunoblot analyses were preformed using antibodies to detect the indicated proteins. Cellular β-actin serves as a loading control.

*ORF50* encodes RTA, an essential immediate-early gene product that is required for early viral gene expression and ultimately viral replication. We therefore evaluated whether *ORF50* deletion correlated with a loss of viral protein production and inhibition of viral replication. Consistent with *ORF50* deletion and the requirement for RTA in promoting viral gene expression, MHV68 proteins were not detected in immunoblot analysis using MHV68 antiserum on lysates from 3T3 cells expressing inducible-Cre (3T3-iCre) infected with O50.loxP (**Fig. 1C**). In contrast, MHV68 antigens were readily detected in lysates from normal 3T3-iCre cells infected with O50.loxP or WT MHV68. Expression of the immediate-early protein LANA was not affected by RTA ablation. In agreement with immunoblot findings, while O50.loxP replicated efficiently in Vero cells, this virus failed to replicate in Vero-Cre cells, while WT MHV68 replicated efficiently in both cell types (**Fig. 2A**). Together, these data demonstrate that Cre expression leads to efficient deletion of floxed *ORF50* and thereby inhibits downstream production of viral proteins and ultimately viral replication. Moreover, the presence of *loxP* sites flanking *ORF50* did not adversely impact progression through the lytic replication cycle.

**Fig 2.**
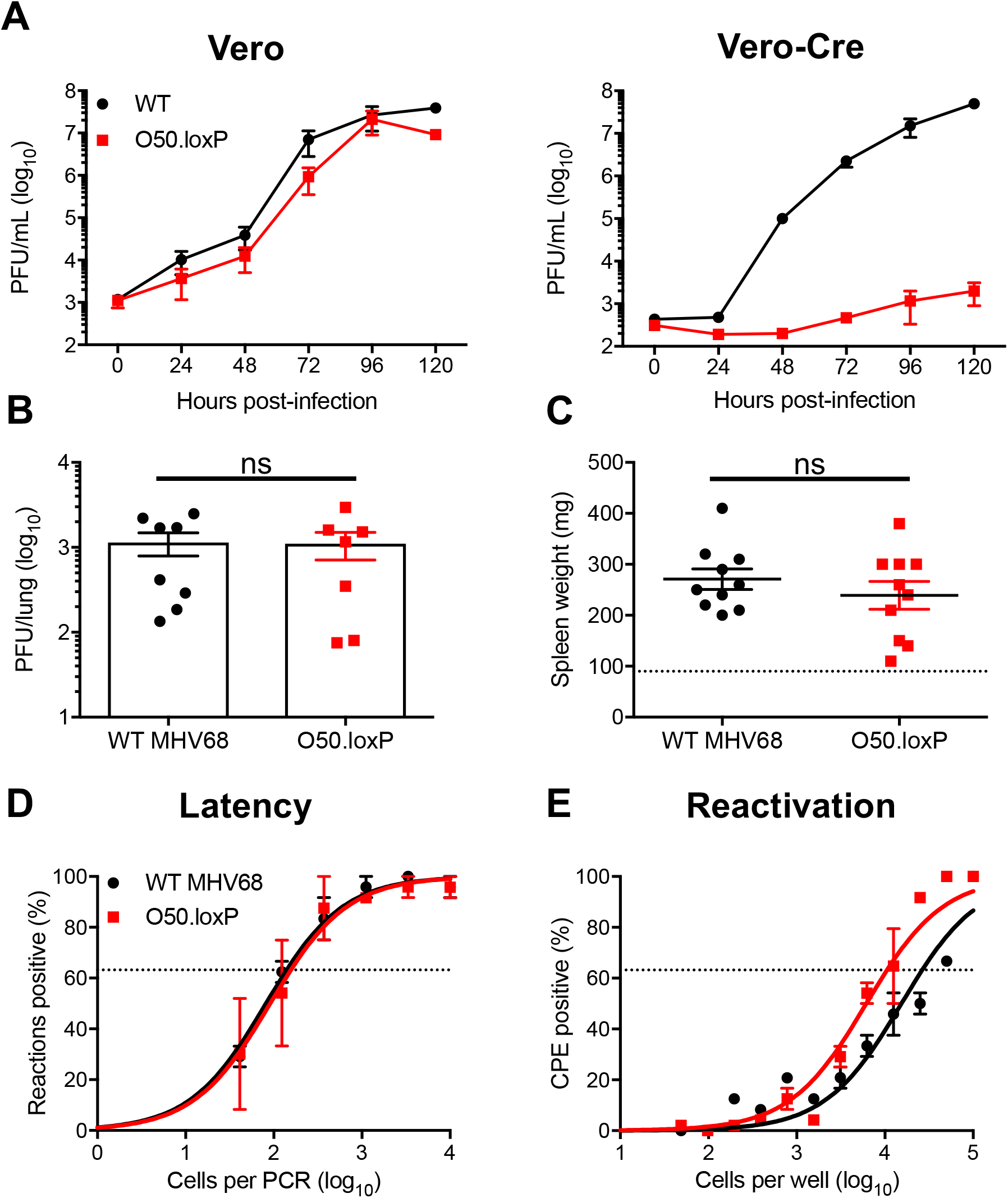
O50.loxP exhibits efficient acute replication, latency, and reactivation in C57BL/6 mice. (A) Vero cells and Vero-Cre cells were infected with WT MHV68 or O50.loxP MHV68 at 0.05 PFU/cell. Viral titers were determined by plaque assay at the indicated times post-infection. Results are means of triplicate samples. Error bars represent standard deviations. (B to E) C57BL/6 mice were infected IN with 1000 PFU of the indicated viruses. (B) Mice were sacrificed on day 7 post-infection, and viral titers in lung homogenates were determined by plaque assay. Each dot represents one mouse. Error bars represent standard error of the means. ns denotes p > 0.05 in a two-tailed Student’s t-test. (C to E) Mice were sacrificed on days 16-18 post-infection and spleens were collected. (C) Spleens were weighed as a measure of splenomegaly. Each dot represents one mouse. The dashed line indicates the average mass of spleens from mock-infected mice. Each dot represents one mouse. Error bars represent standard error of the means. ns denotes p > 0.05 in a two-tailed Student’s t-test. (D) Single-cell suspensions of spleen cells were serially diluted, and the frequencies of cells harboring MHV68 genomes were determined using a limiting-dilution PCR analysis. (E) Reactivation frequencies were determined by *ex vivo* plating of serially diluted cells on an indicator monolayer. Cytopathic effect was scored 2 to 3 weeks post-plating. Groups of 3 to 5 mice were pooled for each infection and analysis. Results are means of 2 to 3 independent infections. Error bars represent the standard error of the means.

Having validated O50.loxP in cultured cells, we next sought to determine if insertion of *loxP* sites flanking *ORF50* impacts MHV68 lytic replication, latency, and reactivation *in vivo*. We first evaluated the lytic replication capacity of O50.loxP MHV68 relative to WT virus following IN inoculation of WT C57BL/6 mice. On day 7 post-infection, titers of both WT MHV68 and O50.loxP were equivalent in the lungs, indicating that O50.loxP lytic infection is not attenuated *in vivo* (**Fig. 2B**). Evaluating latency establishment and reactivation efficiency in spleens on days 16-18 after IN inoculation using limiting-dilution PCR analyses and CPE assays, respectively, we found that O50.loxP established latency in and reactivated from splenocytes as efficiently as WT MHV68 following infection of C57BL/6 mice (**Fig. 2D** and **2E**; **Table 1 and 2**). These findings demonstrate that the presence of *loxP* sites flanking *ORF50* is not detrimental to latency and reactivation. Spleen weights, a correlate of virus-induced splenomegaly (53), also were comparable between WT MHV68 and O50.loxP infections (**Fig 2C**). Together these data indicate that insertion of *loxP* sites flanking the *ORF50* gene does not attenuate MHV68 infection of mice in the absence of Cre-recombinase.

**Table 1.** Frequency of MHV68 latently infected cells

| Mouse strain | Virus | Route of infection | Cell population | Day post-infection | Latency frequency |
| --- | --- | --- | --- | --- | --- |
| C57BL/6 | WT MHV68 | IN | Spleen | 16 | 1/140 |
|  | 50.loxP | IN | Spleen | 16 | 1/75 |
| CD19-Cre | WT MHV68 | IN | Spleen | 16 | 1/65 |
|  | 50.loxP | IN | Spleen | 16 | 1/100 |
|  | WT MHV68 | IN | Spleen | 42 | 1/1,900 |
|  | 50.loxP | IN | Spleen | 42 | 1/1,700 |
|  | WT MHV68 | IP | Spleen | 16 | 1/140 |
|  | 50.loxP | IP | Spleen | 16 | 1/550 |
|  | WT MHV68 | IP | PEC | 16 | 1/500 |
|  | 50.loxP | IP | PEC | 16 | 1/575 |

**Table 2.** Frequency of reactivation competent cells

| Mouse strain | Virus | Route of infection | Cell population | Day post-infection | Reactivation frequency<sup>a</sup> |
| --- | --- | --- | --- | --- | --- |
| C57BL/6 | WT MHV68 | IN | Spleen | 16 | 1/25,000 |
|  | 50.loxP | IN | Spleen | 16 | 1/11,000 |
| CD19-Cre | WT MHV68 | IN | Spleen | 16 | 1/7,800 |
|  | 50.loxP | IN | Spleen | 16 | BLD |
|  | WT MHV68 | IP | Spleen | 16 | 1/4,500 |
|  | 50.loxP | IP | Spleen | 16 | BLD |
|  | WT MHV68 | IP | PEC | 16 | 1/1200 |
|  | 50.loxP | IP | PEC | 16 | 1/2000 |

<sup>a</sup>BLD, below the limit of detection of 1 in 100,000 cells.

### *Deletion of* ORF50 *in B cells does not prevent MHV68 latency establishment*

To test the hypothesis that productive viral replication in B cells influences MHV68 latency, we infected mice that encode Cre-recombinase under the control of the CD19 pan-B cell promoter (54), to allow for B cell-specific deletion of *ORF50* in O50.loxP infected animals. On day 7 post-infection, O50.loxP titers were modestly lower than WT virus titers in lungs of CD19-Cre mice (**Fig. 3A**). Compared to WT MHV68 infection, splenomegaly was significantly reduced in CD19-Cre mice infected with O50.loxP (**Fig. 3B**), suggesting that lytic replication in B cells contributes to the MHV68 IM-like syndrome. However, O50.loxP established latency in spleens of CD19-Cre mice at levels equivalent to WT MHV68 (**Fig. 3C**; **Table 1**). Consistent with the requirement for RTA in GHV replication, reactivation by O50.loxP was severely compromised in CD19-Cre mice (**Fig. 3D**; **Table 2**). Together, these data suggest that the presence of RTA in B cells is critical for MHV68 reactivation from splenocytes and for infection-associated splenomegaly, but is not required for latency establishment in the spleen following IN inoculation.

**Fig 3.**
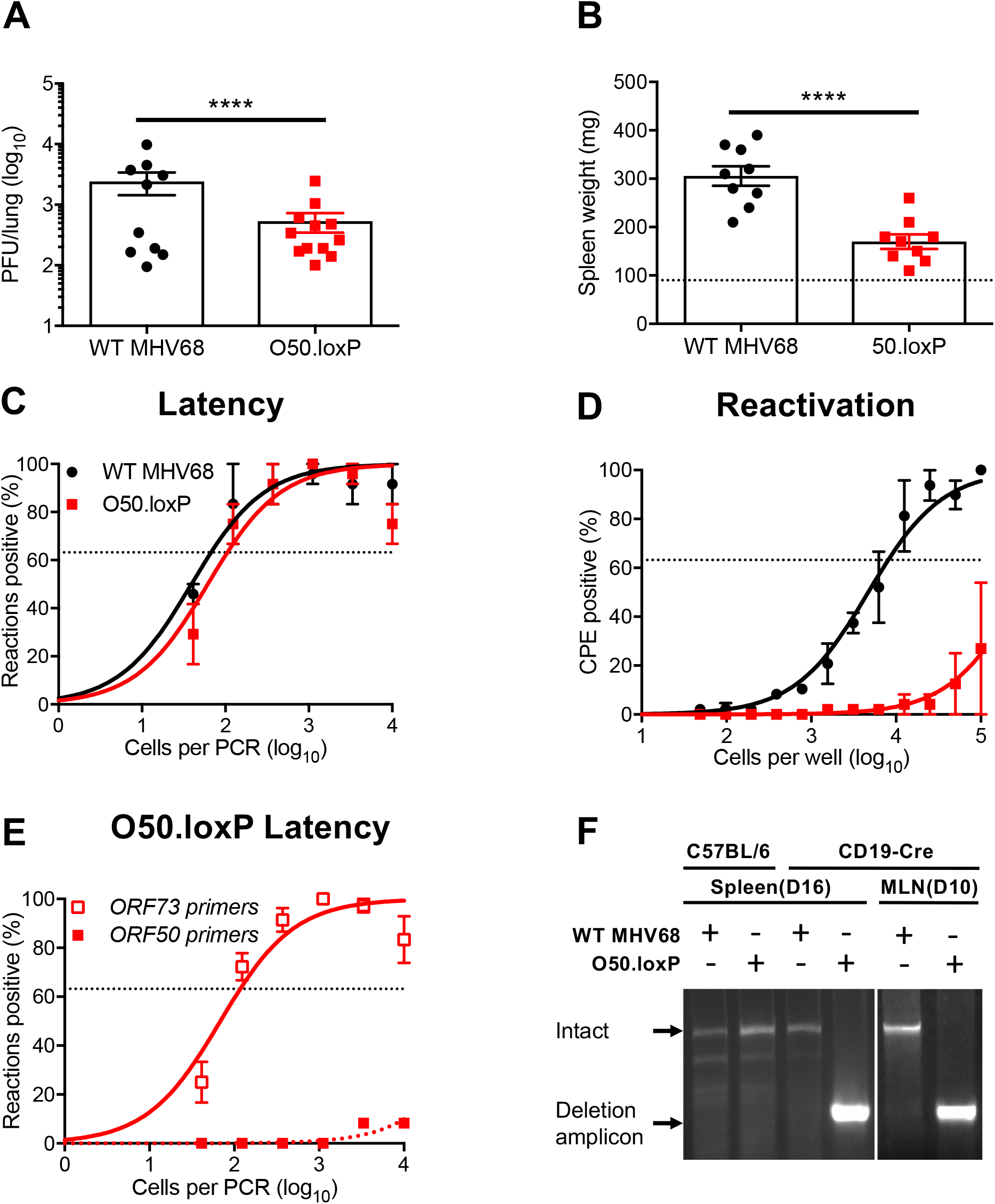
O50.loxP replicates efficiently and establishes latency, but reactivation is impaired in CD19-Cre mice. CD19-Cre mice were infected IN with 1000 PFU of the indicated viruses. (A) Mice were sacrificed on day 7 post-infection, and viral titers in lung homogenates were determined by plaque assay. (B to E) Mice were sacrificed on day 16 post-infection. (B) Spleens were harvested and weighed as a measure of splenomegaly. The dashed line indicates the average mass of spleens from mock-infected mice. Each dot represents one mouse. (C) The frequency of cells harboring MHV68 genomes was determined by limiting-dilution PCR analysis. (D) Reactivation frequencies were determined by *ex vivo* plating of serially diluted cells on an indicator monolayer. Cytopathic effect was scored 2 to 3 weeks post-plating. (E) *ORF50* deletion was confirmed by comparing limiting-dilution PCR analyses performed using primers specific for either the *ORF50* locus or *ORF73* locus. (F) Total DNA was isolated from spleens or lymph nodes of infected C57BL/6 or CD19-Cre mice on days 10 or 16 post-infection with the indicated virus. PCR was performed to evaluate the integrity of the *ORF50* locus. Groups of 3 to 5 mice were pooled for each infection and analysis. Results are means of 2 to 3 independent infections. Error bars represent standard error of the means. **** denotes *P* < 0.0001 in a two-tailed Student’s *t* test.

To confirm that *ORF50* was efficiently deleted following O50.loxP infection of CD19-Cre mice, we performed a parallel LD-PCR analysis of latently infected splenocytes using primers specific to *ORF50* (55), which should be deleted from the viral genome and therefore undetectable (**Fig. 3E**). Although frequencies of splenocytes harboring WT MHV68 genomes were identical when comparing *ORF73* to *ORF50* primers (33), O50.loxP MHV68 genomes were essentially undetectable with *ORF50*-specific primers in CD19-Cre mice (**Fig. 3E**). As an additional control to confirm deletion of *ORF50* from CD19-Cre B cells, we isolated DNA from the mediastinal lymph nodes (MLNs) and spleens of CD19-Cre mice. Using the same PCR approach designed to validate *ORF50* deletion in cultured cells (see **Fig. 1**), only the deletion amplicon was detected from MLNs and spleens of CD19-Cre mice infected with O50.loxP MHV68 (**Fig. 3F**). Since several lines of evidence indicate that MHV68 passes through draining lymph nodes prior to systemic infection (33, 35–37), these observations also suggest that *ORF50* deletion occurs early during the MHV68 dissemination process in CD19-Cre mice. These controls therefore further support that lytic replication in B cells is not a pre-requisite for MHV68 to disseminate to the spleen and establish latency following IN inoculation of mice.

In addition to B cells, peritoneal macrophages also harbor latent MHV68, especially following intraperitoneal (IP) inoculation. To gain a more complete understanding of how *ORF50* deletion in B cells influences MHV68 colonization of the host, we assessed latency and reactivation following IP inoculation of CD19-Cre mice. Consistent with results following IN infection, O50.loxP MHV68 established latency at frequencies comparable to WT MHV68 in splenocytes, while reactivation was substantially reduced (**Fig. 4A and 4B; Table 1 and 2**). In peritoneal exudate cells (PECs), O50.loxP established latency in PECs of CD19-Cre mice at levels equivalent to WT MHV68 and did not exhibit a deficit in *ex vivo* reactivation, which agrees with the notion that macrophages, which are CD19^-^ and should not express Cre-recombinase in CD19-Cre mice, are the major cell type in the peritoneal cavity infected by MHV68 (**Fig. 4C and 4D; Table 1 and 2**). These findings support the conclusion that reactivation from splenocytes, but not PECs, primarily involves B cells, and further suggests that MHV68 does not pass through a B cell prior to infecting the majority of PECs, following IP infection of mice.

**Fig 4.**
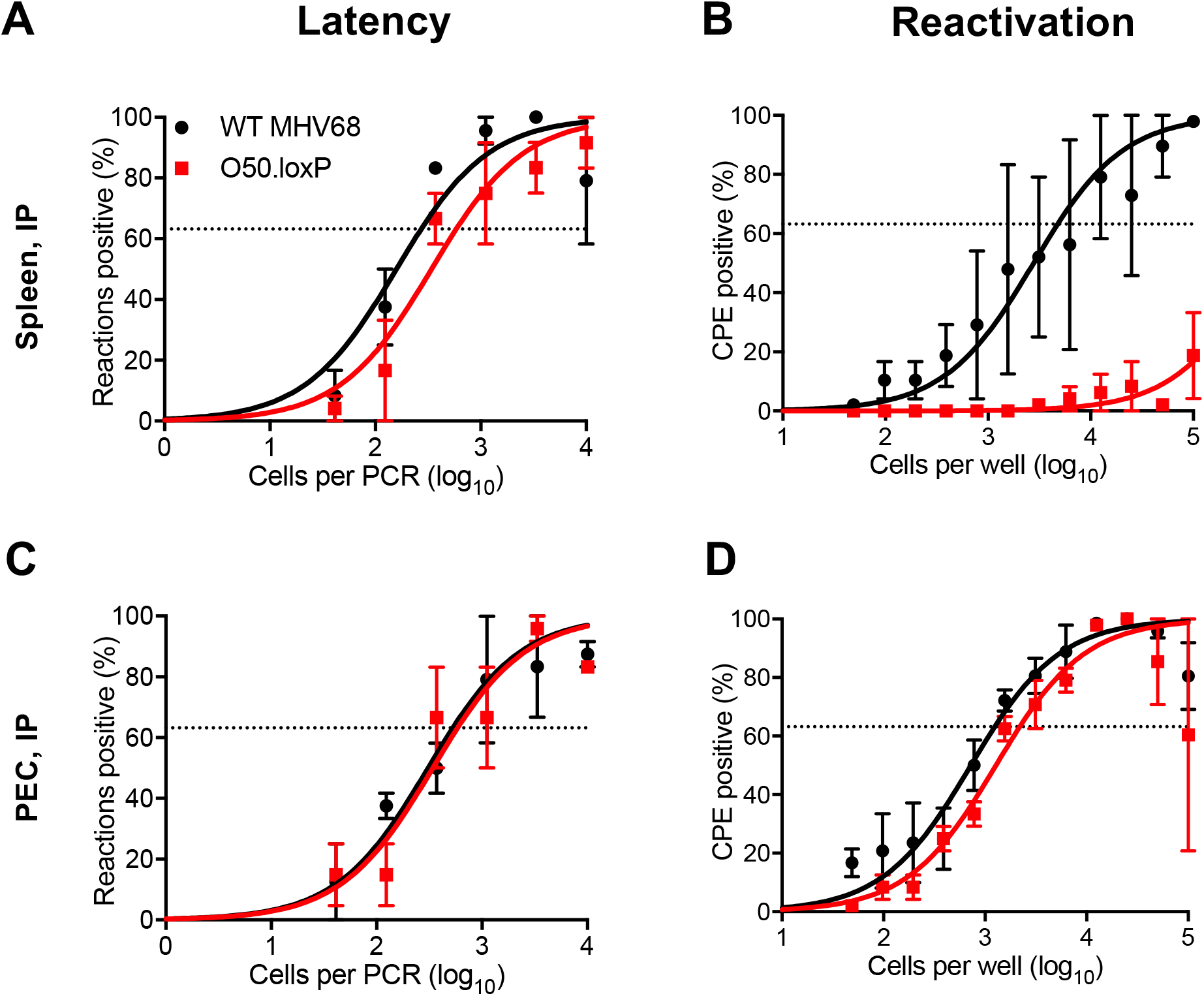
O50.loxP establishes latency in spleens of CD19-Cre mice, but does not reactivate, after IP infection. CD19-Cre mice were infected IP with 1000 PFU of the indicated viruses. Mice were sacrificed on day 16-18 post-infection and spleens (A and B) or peritoneal exudate cells (PECs, C and D) were isolated. (A and C) The frequencies of cells harboring latent viral genomes was determined by limiting-dilution PCR analysis. (B and D) Reactivation frequencies were determined by *ex vivo* plating of serially diluted cells on an indicator monolayer. Cytopathic effect was scored 2 to 3 weeks post-plating. Groups of 3 to 5 mice were pooled for each infection and analysis. Results are means of 2 to 3 independent infections. Error bars represent standard error of the means.

### Proliferation of infected B cells is sufficient for maintaining latent reservoirs

It stands to reason that reactivating virus is a major contributor to primary GHV infections of new hosts. However, whether periodic reactivation occurs within a host already colonized by a GHV in a manner that contributes to the homeostatic maintenance of latent viral reservoirs is not yet clear. Toll-like receptor (TLR) agonists stimulate GHV reactivation *ex vivo* and promote expansion of the latent MHV68 reservoir *in vivo* when administered to mice (48, 49). This supports the hypothesis that reactivation facilitates reseeding of the latent pool. However, it is also possible that B cell proliferation driven by immune activation enables re-expansion of the latent reservoir without overt reactivation – a hypothesis that has not been directly tested.

To begin addressing this question, we first evaluated O50.loxP maintenance over time relative to WT MHV68 in CD19-Cre mice. By day 42 post-infection, a time point at which lytic viral replication has cleared and immune activation has waned (19, 32, 56), we found that *ORF50*-deleted MHV68 was present at levels roughly equivalent to WT virus (**Fig. 5A and Table 1**). Control LD-PCR performed using primers specific to *ORF50* indicated that the locus was in fact absent from viral genomes in splenocytes (**Fig. 5B**), which demonstrates that non-deleted WT virus has not expanded to reclaim the available space. While these observations do not directly address whether homeostatic reactivation is needed to seed naïve cells in order to maintain a latency reservoir, they do indicate that O50.loxP MHV68 maintains latency essentially as well as WT virus in CD19-Cre splenocytes despite a severe defect in reactivation from B cells.

**Fig 5.**
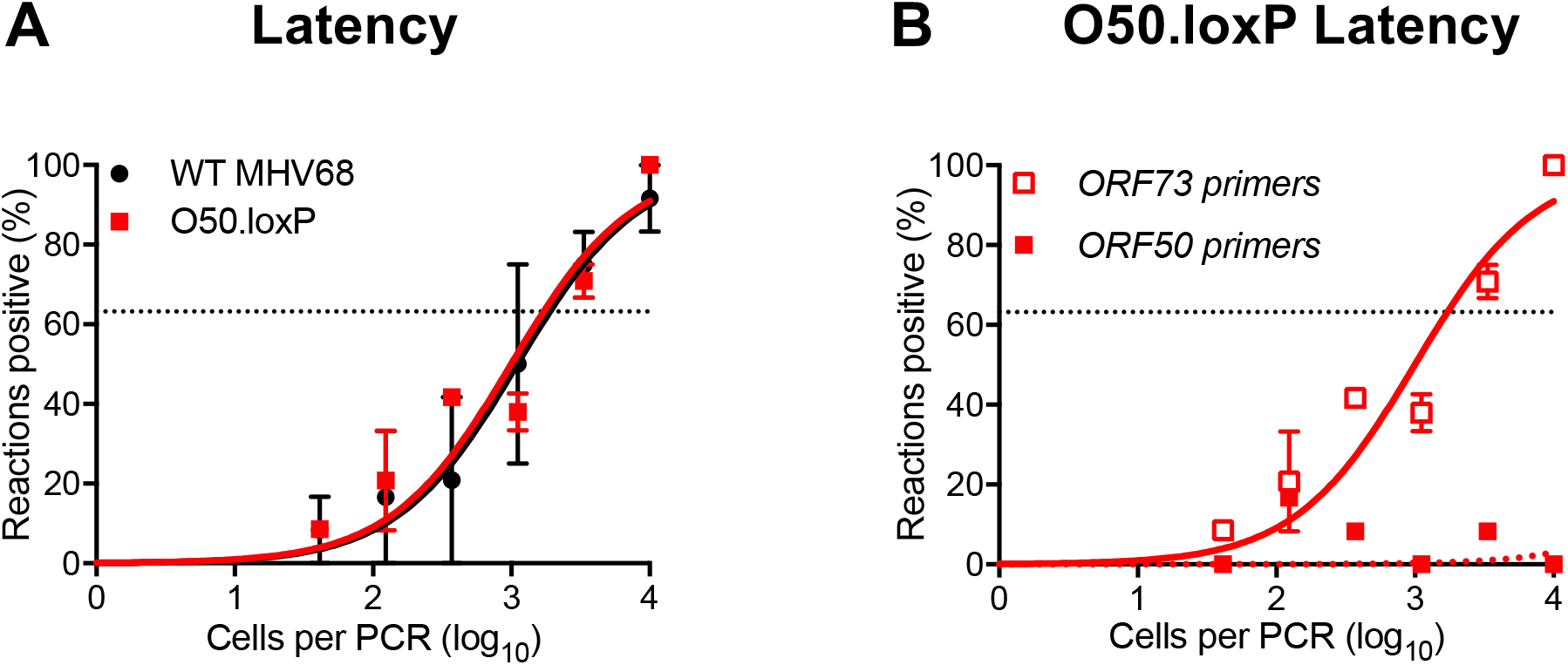
O50.loxP latency is maintained over time despite *ORF50* deletion. CD19-Cre mice were infected IN with 1000 PFU of the indicated viruses. Mice were sacrificed on day 42 post-infection, and spleens were collected. (A) The frequency of cells harboring MHV68 genomes was determined by limiting-dilution PCR analysis. (B) *ORF50* deletion was confirmed by comparing limiting-dilution PCR analyses performed using primers specific for either the *ORF50* locus or *ORF73* locus. Groups of 3 to 5 mice were pooled for each infection and analysis. Results are means of 2 to 3 independent infections. Error bars represent standard error of the means.

We previously demonstrated that lipopolysaccharide (LPS), a TLR4 agonist that stimulates polyclonal B cell activation in mammals, drives MHV68 reactivation from latency *ex vivo* and promotes re-expansion of the MHV68 latency reservoir in spleens of mice (48). Viral recrudescence in lungs and a correlating virus-specific CD8^+^ T cell response support the hypothesis that reactivation promoted the increased latency observed in these experiments (48), but it also is possible that cellular proliferation following LPS treatment contributed to the observed increase in MHV68 latency. Thus, in order to push the conditional deletion system and more definitively test whether proliferation of latently infected B cells is a major contributor to maintaining latency reservoirs, we treated latently infected mice with LPS on day 42 post-infection (See schematic in **Fig. 6A**). In LD-PCR analyses performed two weeks after *in vivo* stimulation with LPS, both O50.loxP and WT MHV68 exhibited 5-10-fold increases in the number of latently infected cells per spleen (1 in 2000 vs. 1 in 1000 cells viral genome-positive, respectively) relative to mock-treated animals at the same time point (1 in ~10^4^ cells for both viruses) (**Fig. 6B** and **Table 3**). LD-PCR performed with primers specific to *ORF50* indicated that the latent viral genomes present did not contain WT viral genomes in which *ORF50* was not deleted (**Fig. 6C**). EdU labeling of cells actively replicating DNA after treatment confirmed that LPS treatment effectively induced cellular proliferation in the spleen (**Fig. 6D**). Given that O50.loxP and WT latency similarly expanded after treatment, these data suggest that LPS drives the re-expansion of latency *in vivo* through cellular proliferation in the absence of MHV68 reactivation from B cells.

**Fig 6.**
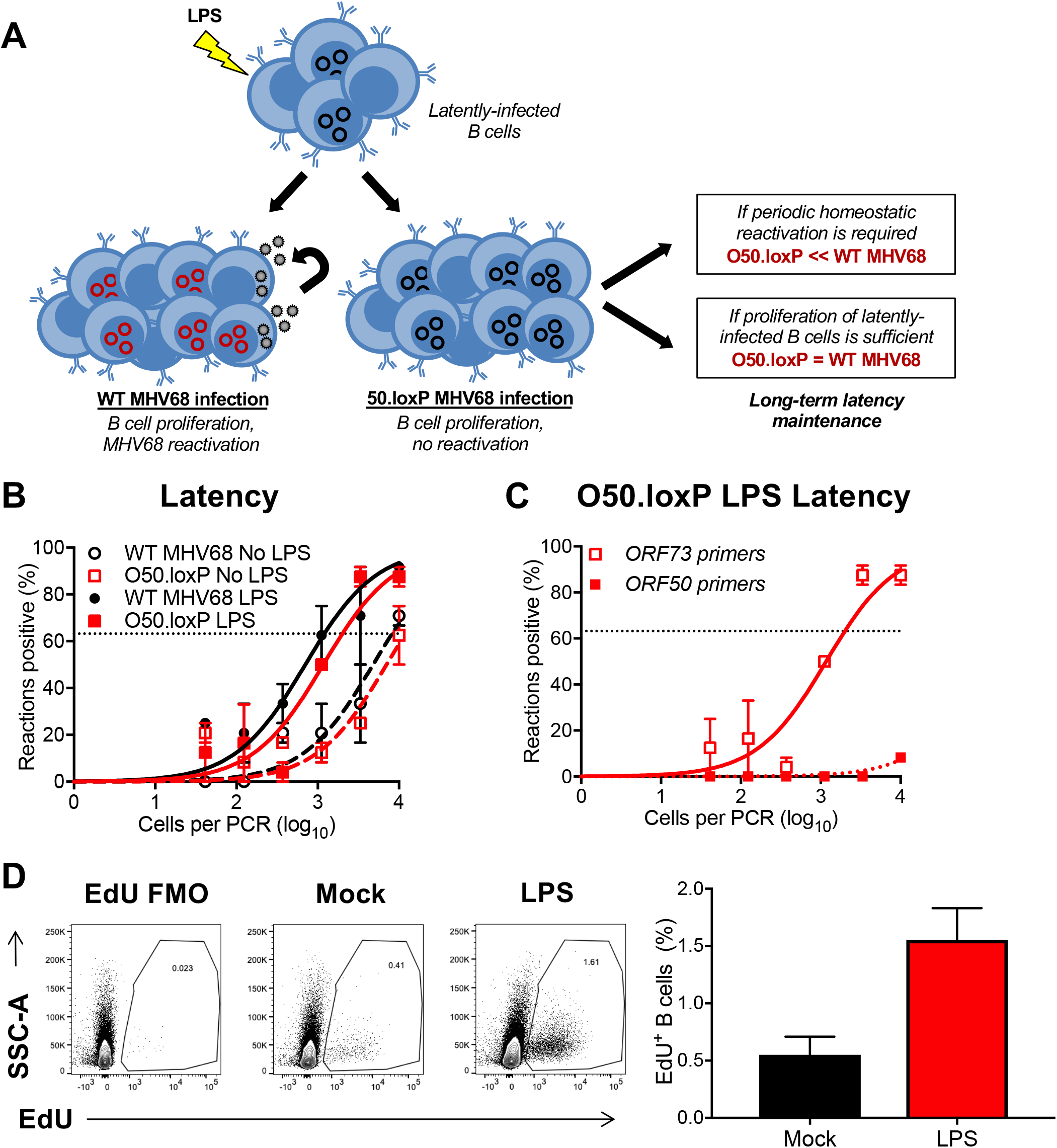
Stimulating B cell proliferation increases O50.loxP genome frequencies. (A) Hypothesized outcomes following polyclonal B cell activation in the presence or absence of MHV68 reactivation from B cells. (B) CD19-Cre mice were infected IN with 1000 PFU of the indicated viruses. On day 42 post-infection mice were injected intraperitoneally with diluent (PBS) or 15 μg of LPS. Mice were sacrificed 14 days post-treatment (56 dpi), and spleens were harvested. (B) The frequency of cells harboring MHV68 genomes was determined by limiting-dilution PCR analysis. (C) *ORF50* deletion was confirmed by comparing limiting-dilution PCR analyses performed using primers specific for either the *ORF50* locus or *ORF73* locus. The LPS-treated O50.loxP infection is shown. Groups of 3 to 5 mice were pooled for each infection and analysis. Results are means of 2 to 3 independent infections. Error bars represent standard error of the means. (D) To confirm that LPS stimulated B cell proliferation, an intraperitoneal injection of EdU was performed 3.5 h prior to sacrifice. Spleen cells with EdU incorporation in newly synthesized DNA were labeled with Click-IT chemistry. EdU^+^ B cells were detected by flow cytometry. B cells were gated as CD19^+^/B220^+^.

**Table 3.** Frequency of MHV68 latently infected cells following LPS treatment

| Mouse strain | Virus | Route of infection | Cell population | Day post-infection | Treatment | Latency frequency |
| --- | --- | --- | --- | --- | --- | --- |
| CD19-Cre | WT MHV68 | IN | Spleen | 58 | No LPS | 1/8,100 |
|  | 50.loxP | IN | Spleen | 58 | No LPS | 1/11,000 |
|  | WT MHV68 | IN | Spleen | 58 | LPS | 1/1,200 |
|  | 50.loxP | IN | Spleen | 58 | LPS | 1/2,000 |

Although reactivation from B cells, the major long-term reservoir for MHV68, does not appear to influence re-expansion of the latent reservoir, a potential caveat to this interpretation is that latent O50.loxP MHV68 within non-Cre-expressing CD19^-^ cells, such as macrophages, dendritic cells, or epithelial cells, could hypothetically reactivate and reseed latency in the spleen. To address this possibility, latently-infected mice were treated with the antiviral drug cidofovir over a 15-day dosing regimen beginning one day prior to LPS administration (**Fig. 7A**). Cidofovir is an inhibitor of the viral DNA polymerase that potently inhibits MHV68 reactivation and associated diseases *in vivo*, especially using the regimen described (57, 58). Two weeks after stimulation, mice treated with LPS and cidofovir exhibited latency levels for both O50.loxP and WT MHV68 that were on par with levels detected in mice that had received LPS alone (**Fig. 7B and Table 4**). We confirmed in parallel experiments that the cidofovir used effectively blocked MHV68 replication (**Fig. 7C**). These data indicate that reactivation occurring after LPS treatment is not a major contributor to the observed expansion of the latency reservoir *in vivo*. Taken together, these findings support the conclusion that proliferation of latently infected B cells, rather than reactivation *per se*, is sufficient for expanding and maintaining long-term latency within a GHV infected organism.

**Fig 7.**
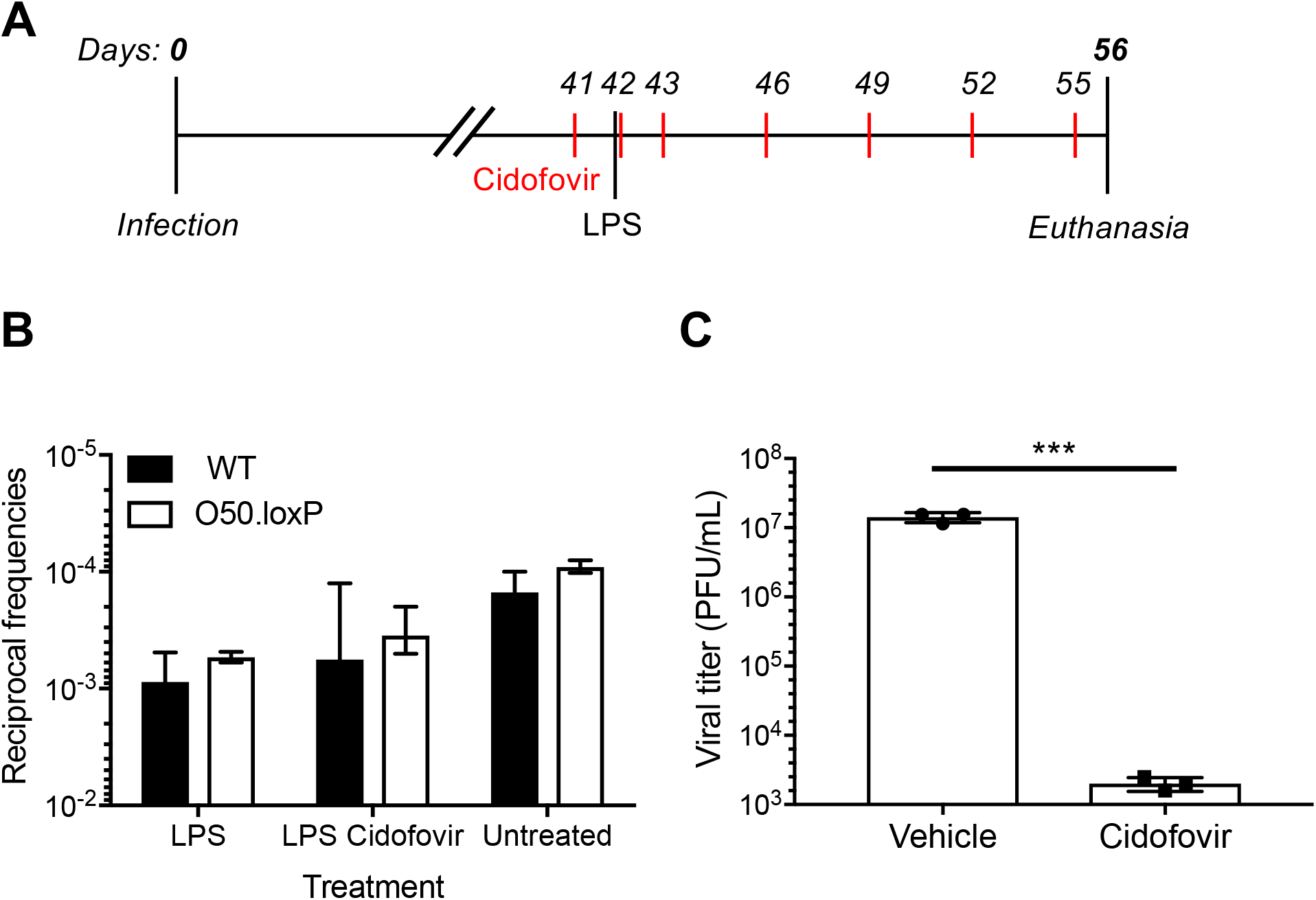
Proliferation of latently infected B cells in the absence of viral reactivation is sufficient for latency maintenance. (A) Experimental design depicting the LPS and antiviral treatment regimens. (B) CD19-Cre mice were infected IN with 1000 PFU of the indicated viruses. Mice were injected subcutaneously with Cidofovir on days 41, 42, 43 and every 3 days subsequently until sacrifice. On day 42 post-infection mice were injected intraperitoneally with diluent (PBS) or 15 μg of LPS. Mice were sacrificed 14 days post-treatment (56 dpi), and spleens were harvested. (B) The frequency of cells harboring MHV68 genomes was determined by limiting-dilution PCR analysis. The reciprocal frequencies of genome positive frequencies were plotted into columns. Results are means of 3 independent infections. (C) To confirm Cidofovir potency, BHK21 cells were treated with vehicle (PBS) or cidofovir (50 ug/mL) prior to infection at an MOI of 5 PFU/cell. Cells were subjected to freeze-thaw lysis 24 h post-infection. Viral titers were determined by plaque assay. Error bars represent standard deviations. *** p < 0.001 in a two-tailed Student’s *t* test.

**Table 4.** Frequency of MHV68 latently infected cells following LPS + Cidofovir treatment

| Mouse strain | Virus | Route of infection | Cell population | Day post-infection | Treatment | Latency frequency |
| --- | --- | --- | --- | --- | --- | --- |
| CD19-Cre | WT MHV68 | IN | Spleen | 58 | No LPS | 1/7,500 |
|  | 50.loxP | IN | Spleen | 58 | No LPS | 1/7,550 |
|  | WT MHV68 | IN | Spleen | 58 | LPS | 1/1,900 |
|  | 50.loxP | IN | Spleen | 58 | LPS | 1/1,950 |
|  | WT MHV68 | IN | Spleen | 58 | LPS +<br>Cidofovir | 1/4,500 |
|  | 50.loxP | IN | Spleen | 58 | LPS +<br>Cidofovir | 1/3,500 |

## DISCUSSION

It is well established that gammaherpesviruses preferentially establish long-term latency in B cells, a cell type that provides a latency reservoir with the capacity to proliferate and survive for the lifetime of the host (18, 25, 40, 54, 59–65). However, whether lytic replication and reactivation from infected B cells contribute to expansion and maintenance of latency is difficult to evaluate with traditional loss-of-function viral mutagenesis strategies. Here we report the use of cell-type-specific, Cre-lox-mediated deletion of *ORF50*, a gene that is essential for viral replication, to examine the importance of lytic replication in B cells in MHV68 latency. The conditional deletion of *ORF50* differs from traditional loss-of-function *ORF50* mutants which are replication-dead in all tissues from the start of the infection process. From our B cell-specific deletion approach, we draw two major conclusions: 1) MHV68 dissemination to and latency establishment in the spleen do not require lytic replication within the B cell compartment, and 2) polyclonal proliferation of latently infected B cells in the absence of viral reactivation is sufficient to maintain a high latent viral load. With this new information, we present a refined model for MHV68 interactions with B cells in the establishment and maintenance of life-long latent infection (**Fig. 8**).

**Fig 8.**
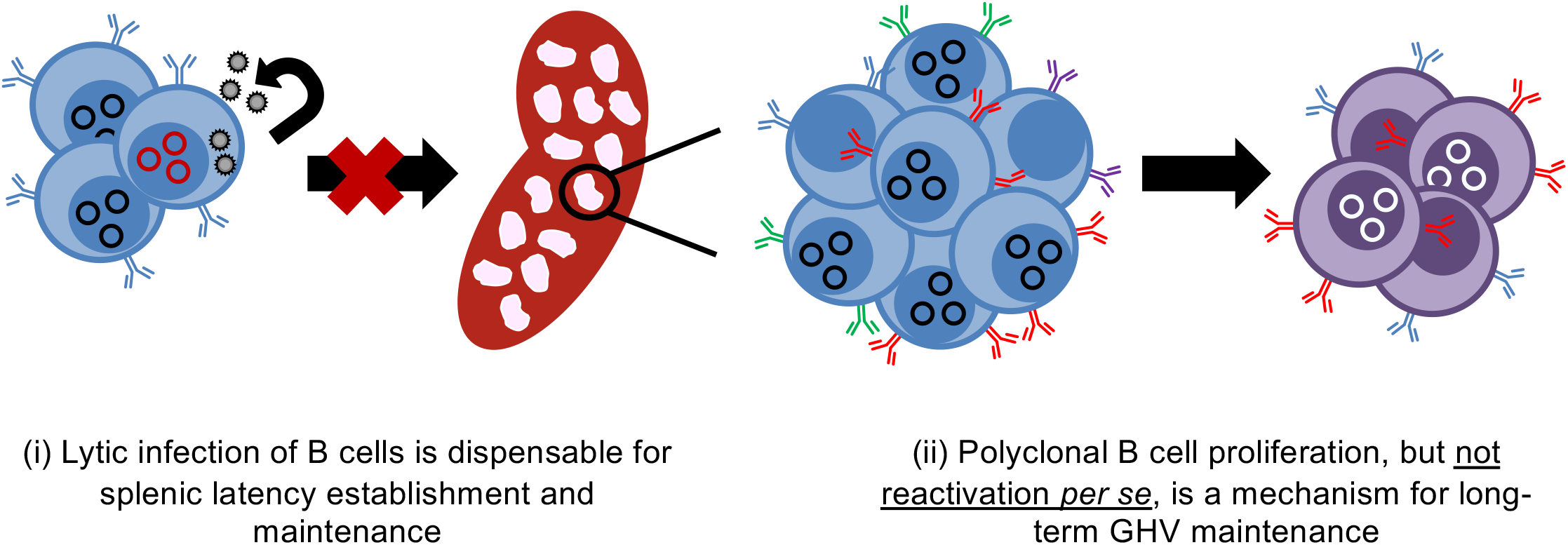
A revised model of chronic MHV68 infection. (i) After lytic infection of the lung epithelium, antigen presenting cells and B lymphocytes traffic the virus to the draining lymph nodes. Lytic replication in B cells is not required for systemic dissemination to distal lymphoid organs, as latency is efficiently established in the absence of *ORF50*. (ii) During long-term chronic infection, B cell proliferation, perhaps in response to other infections, is sufficient to maintain high levels of latently infected B cells. Homeostatic reactivation is not an absolute requirement for maintaining viral genomes within the B cell compartment.

### Viral replication in host colonization

The work presented here sheds new light on requirements for viral dissemination in the host. Following IN inoculation of mice, MHV68 undergoes lytic replication in the respiratory epithelium (32). This promotes infection of lung-resident and recruited antigen presenting cells that carry the virus to draining mediastinal lymph nodes (36, 38). In the draining lymph nodes, the virus is harbored in proliferating B cells before disseminating via hematogenous routes to the spleen (33, 35, 59). Previous work using *ORF50*-null MHV68 or cidofovir treatment to block acute replication following WT MHV68 infection, suggest that lytic replication after IN inoculation is critical for MHV68 dissemination and latency establishment in the spleen (43, 45, 57, 58). In contrast, lung-resident B cells seemingly take up and maintain MHV68 genomes (45). Since B cells facilitate trafficking from lungs to spleens (66) and *ORF50* deletion was evident in draining lymph nodes of CD19-Cre mice by day 10 post-infection with O50.loxP, our data suggest that lytic replication facilitates seeding of B cells in the draining lymph node. This event appears to be sufficient for latent colonization of B cells throughout the host.

Compared to other MHV68 genes we have conditionally deleted in B cells, the observation that O50.loxP MHV68 establishes latency in the spleen at WT levels after IN inoculation is unique. Deletion of the *M2* gene in both CD19^+^ cells and AID^+^ B cells resulted in a ca. 10-fold reduction in splenic latency (42), while B cell-specific deletion of *ORF73*, which encodes the episome maintenance protein LANA, resulted in splenic latency below limits of detection (33). Similar to O50.loxP, however, both M2.loxP and O73.loxP viruses accumulate to levels similar to WT virus in MLNs after IN inoculation and in the spleen after IP infection. This highlights the importance of M2 and LANA functions in B cells in facilitating dissemination to distal sites of latent viral persistence, suggesting that proteins that function in latency are critically important. However, since similar defects in splenic colonization after IN inoculation are not observed for O50.loxP upon B cell-specific deletion of *ORF50*, the M2 and LANA phenotypes further emphasize the conclusion that lytic replication and/or reactivation from B cells does not play a major role in systemic latent infection.

### Contribution of periodic reactivation to latent persistence

With respect to the contribution of reactivation from B cells to long-term latency maintenance, we initially suspected that the frequency of cells harboring O50.loxP MHV68 would more rapidly decay relative to WT reactivation-competent virus. This hypothesis was based on the observation that adoptive transfer of latently infected B cells leads to infection of B cells within a congenic recipient mouse (47) and that TLR agonists that trigger reactivation *ex vivo* also promote increased numbers of latently infected cells when administered to infected mice (48). However, after observing relatively comparable levels of *ORF50*-deleted and WT virus in spleens at later time points, we questioned whether other mechanisms might be at work. Indeed, the finding that LPS, a polyclonal B cell stimulus, promoted essentially equivalent increases in the number of latently infected splenocytes, suggests that GHVs are able to coopt B cell proliferation to maintain high levels of latency. Moreover, the finding that cidofovir did not prevent the LPS driven expansion indicates that increased latency was not due to reactivation of O50.loxP genomes that retained undeleted *ORF50*. These findings strongly suggest that MHV68, and likely EBV and KSHV, uses cellular proliferation, and not reactivation *per se*, as a primary means to maintain B cell latency. This method would enable the viral genome to expand as B cells proliferate in response to other infections, while also limiting the production of antigenic targets that could promote immune detection and clearance of infected cells.

### Comparison to similar studies

Cre-lox recombination provides an elegant system for conditional deletion, but the deletion must be efficient and specific. It is notable that our findings differ from those of a previous study (67), however nuances of the two studies may explain the apparently conflicting results. Contrary to our results, a virus in which both *ORF49* and *ORF50* were floxed (MHV-F50) was significantly attenuated following both IN and IP infection of CD19-Cre mice. The original MHV68 BAC system used *loxP* sites flanking the inserted BAC vector sequences to allow removal of the foreign bacterial origin of replication and selection cassette from the viral genome during virus recovery in Cre-expressing cells (68). To employ Cre-*loxP*-mediated deletion of viral genes, this presents a potential problem, as recombination could occur between a floxed gene of interest and the other *loxP* sites in the genome, leading to deletion of large segments of viral sequences. In a previous study in which the *ORF73* latency gene was conditionally deleted, we described the generation of a new BAC system that employed the Flp-*Frt* recombination system for BAC vector deletion (33). We used this system here to ensure that *ORF50* would not be inadvertently deleted while generating virus stocks. In contrast, the previous MHV-F50 study utilized the original BAC that contains additional *loxP* sites (68). It is possible that this led to a larger than expected deletion within the MHV68 genome in Cre-expressing mice that inadvertently deleted the left end of the viral genome, in addition to *ORF50*.

While we consider the possibility less likely, it also is possible that deletion of both *ORF50* and *ORF49* yields a much more severe attenuated phenotype, than deletion of *ORF50* alone. *ORF49* encodes a protein that is essential for lytic infection, and viruses lacking *ORF49* are attenuated for both lytic and latent infection (69, 70). As a confirmation of our O50.loxP MHV68 data, we also targeted another essential lytic gene, *ORF57*. Similar to O50.loxP, O57.loxP MHV68 disseminated to the spleen and established latency despite deletion of *ORF57* from B cells (Owens, Gupta, and Forrest, manuscript in preparation). This serves as independent confirmation and provides additional confidence in the conclusion that lytic replication in B cells is not necessary for viral dissemination throughout the animal and latency establishment. In conclusion, the data presented here highlight the utility of cell-type specific deletion of viral genes to foster an understanding of fundamental mechanisms of GHV infection and persistence in the host.

## MATERIAL AND METHODS

### Ethics statement

Mouse experiments performed for this study were carried out in accordance with NIH, USDA, and UAMS Division of Laboratory Animal Medicine and Institutional Animal Care and Use Committee (IACUC) guidelines. The protocol supporting this study was approved by the UAMS IUCAC. Mice were anesthetized prior to inoculations and sacrificed humanely at the end of experiments.

### Cells and viruses

Vero (ATCC CCL-81) and Vero-Cre cells, NIH 3T12 (ATCC CCL-164), BHK21 (ATCC CCL-10), and Swiss albino 3T3 (ATCC CRL-1658) fibroblasts were cultured in Dulbecco’s Modified Eagle Medium (DMEM) supplemented with 10% fetal bovine serum (FBS), 100 U/ml penicillin, 100 μg/ml streptomycin, and 2 mM L-glutamine (cDMEM). Cells were cultured at 37°C with 5% CO_2_ and ~99% humidity. Murine embryonic fibroblasts (MEFs) were harvested from C57BL/6 mouse embryos and immortalized as previously described (66). Viruses used in this study included the FRT BAC-derived wild-type MHV68 (henceforth referred to as WT MHV68) (33); the RTA deficient RTA-null MHV68, and the conditional *ORF50* mutant (O50.loxP MHV68) (43, 45). WT and O50.loxP viruses were derived from MHV68 with FRT sites flanking the BAC cassette, and were passaged in Flp-expressing 3T12 cells to remove the BAC cassette. Titers were quantified as described previously.

### Generation of recombinant viruses

The WT MHV68 BAC was created by *en passant* mutagenesis (68). To generate the O50.loxP, *loxP* sites were inserted adjacent to the 5’ and 3’ ends of the *orf50* exon 2 (E2) in a FRT BAC template by two successive rounds of *en passant* mutagenesis (33) utilizing the following primers:

O50loxpUP_for 5’AGTCTGCAAGAAATAATAGCCTCCCACTTTTATGGAAATCATAACTTCGTATAGCATACATTATACGAAGTTATTGGTAGCTCCTCCTACATGATAGGGATAACAGGGTAATCGATT,
O50loxpUP_rev 5’GGTTAATTGGTTGTAACACTGGCCTCCCACTTTTATGGAAATCATAACTTCGTATAGCATACATTATACGAAGTTATTGGTAGCTCCTCCTACATGAGTCATGAAATGTCCCTTCAA,
O50loxpDWN_for 5’CATACTTAGTCCACTCGACCCAAACAGCCTGGAGTCATAAATAACTTCGTATAGCATACATTATACGAAGTTATACGGTGCCAAATACAAGACATTTAGGGATAACAGGGTAATCGATTT,
O50loxpDWN_rev 5’GGTTAATTGGTTGTAACACTGGCCAAACAGCCTGGAGTCATAAATAACTTCGTATAGCATACATTATACGAAGTTATACGGTGCCAAATACAAGACATTTCTAAATCCTTAAGTATAA.

Viruses were passaged in Flp-expressing 3T12 fibroblasts to remove the BAC cassette, and titers were quantified as previously described (33).

### Cell culture infections

For immunoblot assays, 3T12 fibroblasts were infected at a multiplicity of infection (MOI) of 5 PFU/cell with WT MHV68, RTA-null, O50.loxP MHV68. Briefly, viral stocks were diluted in low-volume (200 μl) of cDMEM and added directly onto fibroblast cell monolayers seeded the previous day at a density of 2 × 10^5^ cells / well in 6-well plates. Plates were incubated at 37°C and rocked gently every 15 min for 1 h. After 1 h, cDMEM was added back to the cultures.

For all other viral infections, 3T3 or 3T12 cells plated the previous day were inoculated with a low volume of virus diluted in cDMEM. Plates were rocked every 15 min for 1 h at 37°C. After low-volume adsorption, inocula were removed for high-multiplicity infections. After adsorption, cells were incubated in a normal culture volume of cDMEM for the indicated times at 37°C. For multistep growth curves, infected cells were frozen at −80°C at the indicated time points. Cells were subjected to freeze-thaw lysis to release progeny virions, and lysates were serially diluted for plaque assays as described (71).

### Nucleic acid isolation and PCR

Vero or Vero-Cre cells were mock-infected or infected with MHV68 at a multiplicity of infection (MOI) of 5 PFU/cell. Nucleic acid was isolated from cells at 24 or 48 hours post-infection. Total DNA was isolated using a GenCatch blood and tissue genomic mini-prep kit (Epoch Life Science). PCR for the detection of the full-length *ORF50* gene was performed using primers ORF50_del_for 5’GCTTCCTCGTCTACAGAGGTCAGG and ORF50_del_rev 5’GGCACCCATACTAAGTTGTGATTC. *ORF73* and *GAPDH* genes were detected by PCR utilizing primers 73_IG_US and 73_IG_DS for *ORF73* and GAPDHF and GAPDHR for *GAPDH* in GoTaq polymerase (Promega) as described (33).

### Immunoblot analyses

Immunoblot analyses were performed as previously described (33). Briefly, cells were lysed with radio immunoprecipitation (RIPA) buffer (150 mM NaCl, 20 mM Tris, 2 mM EDTA, 1% NP-40, 0.25% deoxycholate supplemented with phosphatase and protease inhibitors), and protein samples were centrifuged at 16,000×g to remove insoluble debris. Protein content for each sample was quantified using the BioRad DC Protein Assay (BioRad). Samples were diluted in 6X Laemmli sample buffer and resolved by sodium dodecyl sulfate polyacrylamide gel electrophoresis (SDS-PAGE) and transferred to nitrocellulose membranes (Thermo Scientific). Blots were probed with the indicated primary antibodies and with horseradish peroxidase (HRP) conjugated secondary antibodies (Jackson ImmunoResearch). Chemiluminescent signal was detected using a ChemiDoc MP Imaging System (Bio-Rad) on blots treated with Clarity ECL reagent (Bio-Rad).

### Mouse-infections and tissue harvests

Male and female, C57BL/6, heterozygous CD19-Cre [B6.129P2(C)-Cd19tm1(cre)Cgn/J] mice were purchased from Jackson Laboratories or were bred and maintained in the animal housing facilities at the University of Arkansas for Medical Sciences (UAMS, Little Rock, AR). Note: Cre-encoding mice used were heterozygous for Cre expression. Mice were housed in sterile conditions and treated according to the guidelines of the Division of Laboratory Animal Medicine (DLAM) at UAMS. Eight to ten-week old mice were anesthetized using isoflurane and inoculated with 1000 PFU of virus diluted in incomplete DMEM (20 μl) for intranasal inoculations or injected with 1000 PFU of virus diluted in incomplete DMEM (200 μl) for intraperitoneal inoculations. Lungs were harvested 7-10 days post-infection. Serum, splenocytes and peritoneal exudate cells (PECs) were harvested 16 to 18 days post-infection, or 42 and 90-days post-infection, as described previously (55). Cells from draining lymph nodes were isolated as previously described (33).

### Splenocyte isolation and limiting-dilution analyses

Spleens were homogenized using a tenBroek tissue disrupter. Red blood cells were lysed by incubating tissue homogenate in 8.3 g/liter ammonium chloride for 10 minutes at room temperature with shaking. Cells were filtered through 40-μM mesh to reduce clumping. Frequencies of cells harboring MHV68 genomes were determined using limiting-dilution (LD) PCR analysis with primers specific for an *ORF73* target as previously described (33). We also performed the same experiments with primers specific for an *ORF50* gene segment to confirm excision of floxed *ORF50* in infected cells (33). Frequencies of latently infected cells capable of reactivating were determined using a limiting-dilution analysis for cytopathic effect induced on an indicator monolayer as previously described (66).

### Plaque assays

Plaque assays were performed as previously described (71) using BHK21 cells (2 × 10^5^ cells/well). Briefly, infected cells were overlaid with 1.5% methylcellulose in DMEM supplemented with 2.5% calf serum, 100 U/ml penicillin, 100 μg/ml streptomycin, and 2 mM l-glutamine and incubated at 37°C for 4 to 6 days. Cell monolayers were stained with a solution of crystal violet in formalin for identification and quantification of plaques.

### Antiviral treatment and in vivo stimulation of reactivation

For *in vitro* analyses, BHK21 cells were infected at an MOI of 5 PFU/cell. Cells were treated with vehicle (PBS) or cidofovir (50 ug/mL) for 24 hours. Cell lysates were isolated and viral titers were quantified by plaque assay.

For *in vivo* analyses, CD19-Cre mice were injected intraperitoneally with 15 μg of *E. coli* lipopolysaccharide in 100 μl of sterile PBS 42 days post-infection to stimulate a TLR response. Where indicated, mice were subcutaneously injected with 25 mg/kg of cidofovir in sterile PBS on alternating flanks (60-100 μl inoculum volume), on days 41, 42, 43, 46, 49, 52, and 55 post-infection. On day 56 post-infection (day 14 post-LPS, where indicated), mice were euthanized by CO_2_ asphyxiation and spleens were collected for analysis.

For proliferation analysis, mice were injected with 100 μg of 5-ethynyl-2’deoxyuridine (EdU, Invitrogen) 3.5 hours prior to splenocyte harvest. Cells were fixed with 4% paraformaldehyde in PBS and Click-iT chemistry (BD Pharmagen) performed according to the manufacturer’s instructions to fluorescently mark EdU^+^ cells. Blocking and detection antibodies were diluted in phosphate-buffered saline (PBS) with 0.2% bovine serum albumin (BSA) and 1 mM EDTA. Splenocytes were blocked with anti-CD16/32 (BD Biosciences) for 15 minutes before surface staining for 30 minutes in the dark at 4°C. Dead cells were labeled using fixable viability dye eFlour 780 (eBioscience) per the manufacturer’s instructions. Surface stains include dump gate (CD3, Ter119, CD11b, CD11c-PerCp5.5), B220-redFluor 710 (Tonbo), and CD19-BV650 (BioLegend). Cells were analyzed on a BD Fortessa using FACS Diva software. Data were analyzed using FlowJo software (v10.6.2).

### Antibodies and drug treatments

Antibodies used in this study include rabbit polyclonal mLANA anti-serum (72), mouse polyclonal MHV68 anti-serum (71), anti-RTA (73), anti-v-cyclin (74), and mouse monoclonal anti-β-actin (Sigma Aldrich, #A2228). For drug treatments, the 17β-estradiol agonist Z-4-hydroxytamoxifen (4-OHT; Alexis Biochemicals; #ALX-550-361-M001) was dissolved at a stock concentration of 2 mM. Inducible-Cre 3T3 fibroblasts plated the previous day were treated with either 0.2 μM 4-OHT to induce nuclear translocation of Cre or ethanol as a vehicle control for 24 h prior to infection.

### Statistics

All statistical analyses were performed using GraphPad Prism software (GraphPad Software, San Diego, CA). Statistical significance was determined by a two-tailed unpaired Student’s t test with a 95% confidence.

## ACKNOWLEDGMENTS

This work was supported in part by grant R01CA167065 from the NIH National Cancer Institute and funds from the UAMS College of Medicine and Arkansas Biosciences Institute to J.C.F. The Flow Cytometry Core and the work described here were also supported in part by Center for Microbial Pathogenesis and Host Inflammatory Responses award P20GM103625 from the NIH National Institute of General Medical Sciences Centers of Biomedical Research Excellence. S.M.O. was supported by a postdoctoral fellowship from Translational Research Institute (TRI) grant TL1 TR003109 through the NIH National Center for Advancing Translational Sciences. D.W.W. and D.G.O. were supported by the Gundersen Medical Foundation. The funders had no role in study design, data collection and interpretation, or the decision to submit the work for publication.

